# Deletion of the mitochondrial matrix protein cyclophilin-D prevents parvalbumin interneuron dysfunction and cognitive deficits in a mouse model of NMDA hypofunction

**DOI:** 10.1101/2020.04.15.043570

**Authors:** Aarron Phensy, Kathy L. Lindquist, Karen A. Lindquist, Dania Bairuty, Esha Gauba, Lan Guo, Jing Tian, Heng Du, Sven Kroener

**Affiliations:** School of Behavioral and Brain Sciences, The University of Texas at Dallas, Richardson, TX 75080; Department of Biological Sciences, The University of Texas at Dallas, Richardson, TX 75080

## Abstract

Redox dysregulation and oxidative stress are final common pathways in the pathophysiology of a variety of psychiatric disorders, including schizophrenia. Oxidative stress causes dysfunction of GABAergic parvalbumin-positive interneurons (PVI), which are crucial for the coordination of neuronal synchrony during sensory- and cognitive-processing. Mitochondria are the main source of reactive oxygen species (ROS) in neurons and they control synaptic activity through their roles in energy production and intracellular calcium homeostasis. We have previously shown that in male mice transient blockade of NMDA receptors during development (subcutaneous injections of 30 mg/kg ketamine (KET) on postnatal days 7, 9, and 11) results in long-lasting alterations in synaptic transmission and reduced parvalbumin expression in the adult prefrontal cortex (PFC), contributing to a behavioral phenotype that mimics multiple symptoms associated with schizophrenia. These changes correlate with oxidative stress and impaired mitochondrial function in both PVI and pyramidal cells. Here, we show that genetic deletion (*Ppif*^*-/-*^) of the mitochondrial matrix protein cyclophilin D (CypD) prevents perinatal KET-induced increases in ROS and the resulting deficits in PVI function, and changes in excitatory and inhibitory synaptic transmission in the PFC. Deletion of CypD also prevented KET-induced behavioral deficits in cognitive flexibility, social interaction, and novel object recognition. Taken together, these data highlight how mitochondrial activity may play an integral role in modulating PVI-mediated cognitive processes.

**Significance Statement:** Mitochondria are important modulators of oxidative stress and cell function, yet how mitochondrial dysfunction affects cell activity and synaptic transmission in psychiatric illnesses is not well understood. NMDA receptor blockade with ketamine during development causes oxidative stress, dysfunction of parvalbumin-positive interneurons (PVI), and long-lasting physiological and behavioral changes. Here we show that mice deficient for the mitochondrial matrix protein cyclophilin D show robust protection from PVI dysfunction following perinatal NMDAR-blockade. Mitochondria serve as an essential node for a number of stress-induced signaling pathways and our experiments suggest that failure of mitochondrial redox regulation can contribute to PVI dysfunction.

## Introduction

Schizophrenia is a neurodevelopmental disorder in which genetic risk factors and early life stressors converge (Harrison and Owen, 2003). Genes involved in glutamatergic synaptic transmission figure prominently among both the rare (Timms et al., 2013; Pocklington et al., 2015; Sekar et al., 2016) and common (Pers et al., 2016) gene variants that contribute to the heritable risk for schizophrenia. In the frontal cortex, these genes are highly expressed early in development (Gulsuner et al., 2013; Birnbaum et al., 2015). Dysfunction of glutamatergic NMDARs during neurodevelopment can disrupt maturation of interneurons (Zhang and Sun, 2011) and cause abnormalities in the GABAergic and dopaminergic systems in schizophrenia (Olney et al., 1999; Krystal et al., 2002; Catts et al., 2013). Aberrant NMDAR activity can therefore shift the cortical excitation-inhibition (E/I) balance (Insel, 2010; Lewis et al., 2012), leading to increased basal neural activity (Jadi et al., 2016), excessive glutamatergic release (Plitman et al., 2014), and oxidative stress (Hardingham and Do, 2016; Steullet et al., 2016). In support of this, NMDAR antagonists such as phencyclidine and ketamine (KET) induce a schizophrenia-like syndrome in healthy subjects, and exacerbate symptoms in schizophrenic patients (Malhotra et al., 1997; Krystal et al., 2002; Anticevic et al., 2012). Pharmacological blockade or genetic deletion of NMDARs in rodents mimics many of the behavioral symptoms seen in patients with schizophrenia, and it also reduces markers of GABAergic interneurons, including cells that express the calcium-binding protein parvalbumin (Abekawa et al., 2007; Braun et al., 2007; Belforte et al., 2010). Reductions in parvalbumin expression are a core finding of post-mortem studies in schizophrenia patients (Olney et al., 1999; Lewis et al., 2005; Akbarian and Huang, 2006) that is replicated by virtually all animal models of the disease (Jiang et al., 2013; Steullet et al., 2017). Fast-spiking PVI have a unique metabolic profile that is reflected in a large number of mitochondria and enriched cytochrome c oxidase (Kann and Kovacs, 2007), which seems to make them particularly susceptible to external stressors during development (Hardingham and Do, 2016; Steullet et al., 2017).

Mitochondria are crucial regulators of oxidative and nitrosative stress (Chen et al., 2003; Li et al., 2004), and transcriptomic, proteomic, and metabolomic studies in post-mortem samples from subjects with schizophrenia indicate alterations in the expression of several proteins associated with mitochondrial function (Prabakaran et al., 2004; Altar et al., 2005; Iwamoto et al., 2005; Hjelm et al., 2015). Altered levels of ATP and mitochondrial dysfunction in the frontal lobe are correlated with negative symptoms, as well as cognitive and memory deficits in schizophrenia (Ben-Shachar and Laifenfeld, 2004; Rajasekaran et al., 2015). Oxidative and other cellular stresses promote translocation of the mitochondrial matrix protein cyclophilin D (CypD) to the inner membrane. This translocation triggers the opening of the mitochondrial permeability transition pore (mPTP) (Connern and Halestrap, 1994; Baines et al., 2005), which is important in glutamate excitotoxicity that results from overactivation of glutamate receptors and subsequent excessive calcium entry into the cell (Schinder et al., 1996; White and Reynolds, 1996). Prolonged CypD-mediated opening of mPTP causes collapsed mitochondrial membrane potential, elevated mitochondrial ROS generation, and lowered ATP production, leading to metabolic changes and ultimately cell death (Basso et al., 2005; Halestrap, 2010). Because CypD is a necessary component of the mPTP, reducing CypD translocation in order to block mPTP formation can preserve mitochondrial function (Du and Yan, 2010).

Here, we investigated whether genetic deletion of CypD (*Ppif*^*-/-*^) can prevent changes in PVI function, PFC physiology, and behavior that develop in a well-characterized rodent model of NMDA hypofunction. Perinatal treatment with ketamine induced oxidative stress and reduced PV expression in the PFC of wildtype-but not of *Ppif*^*-/-*^ mice. CypD-deletion similarly protected against changes in glutamatergic transmission at PVI and deficits in cognitive flexibility, social interaction, and novel object recognition. These data indicate that mitochondrial redox regulation is an important contributor to PVI dysfunction and the resulting E/I imbalance that results from NMDAR-hypofunction.

## Materials and Methods

### Transgenic *Ppif*^*-/-*^ mice

Cyclophilin-D knockout mice (B6;129-*Ppif*^tm1Jmol^/J; The Jackson Laboratory; RRID:IMSR_JAX:009071), were crossed with G42 mice (CB6-Tg[Gad1-EGFP]G42Zjh/J; The Jackson Laboratory; RRID:IMSR_JAX:007677) which express GFP in PVI neurons, in order to identify PV+ neurons in slice electrophysiology experiments. First-and second-generation breeders were selected and their WT (CB6-TgWT; Gad1-EGFP) and *Ppif*^*-/-*^ (CB6-Tg*Ppif*^*-/-*^; Gad1-EGFP) offspring were used for experiments. All procedures were approved by the Institutional Animal Care and Use Committee of The University of Texas at Dallas.

### Perinatal ketamine treatment

On postnatal day 7, 9, and 11 mice received subcutaneous injections of either saline or a sub-anesthetic dose of the NMDA-antagonist ketamine (30mg/kg; Ketathesia HCL, Henry Schein). Both *Ppif*^*-/-*^ and WT mice received either perinatal ketamine (KET) or saline injections, creating four groups: Wild type mice that received saline injections (WT-SAL); Wild type mice that received ketamine injections (WT-KET); *Ppif*^*-/-*^ mice that received saline injections (*Ppif*^*-/-*^-SAL); and *Ppif*^*-/-*^ mice that received ketamine treatment (*Ppif*^*-/-*^-KET). All experiments were performed on adult male *Ppif*^*-/-*^ and WT mice (60-120 days old). Animals that participated in behavioral experiments were handled for 5 minutes a day in the vivarium for 2 weeks prior to the test and then also in the room in which behavioral testing took place for 3 days prior to the test. On the day of testing, animals were transferred to the behavioral room at least 30 min before testing began. Behavioral testing and analysis was performed by experimenters blind to the experimental condition of the subjects.

### Immunohistochemistry

Animals were perfused transcardially with saline for 2 minutes, followed by 4% paraformaldehyde in 0.12 M phosphate-buffered saline (PBS), at 4°C, pH 7.4; Fisher Scientific) for 10 minutes using a peristaltic pump (5.5 ml/min; PeriStar Pro, WPI). Brains were post-fixated in paraformaldehyde with 30% sucrose for 1 h and then transferred to 30% sucrose in PBS for 18 h at 4°C. Coronal slices (40 um) were cut on a freezing microtome. Free-floating sections were incubated in rabbit anti-parvalbumin (1:2000 working dilution; Swant Cat# PV 25, RRID:AB_10000344) in PBS and 0.3% Triton X (Sigma-Aldrich) for 36 h at 4°C. Sections were washed three times for 10 min each in PBS before they were incubated in secondary 594 goat anti-rabbit (1:1000 working dilution; Jackson ImmunoResearch Labs Cat# 111-585-144, RRID:AB_2307325) in PBS and 0.3% Triton X. Sections were washed, mounted, and cover slipped using Prolong Gold Antifade with DAPI (Thermo Fisher Scientific). To quantify PVI immunofluorescence a minimum of four sections from each animal containing the prelimbic and infralimbic regions of the PFC were imaged on a confocal microscope (FluoView 1000, Olympus) at 20x magnification. The number of PV+ cells were hand counted in ImageJ (National Institutes of Health), and DAPI-labeled cells were counted using the thresholding function in ImageJ to obtain the percentage of total PV+ cells among all DAPI-labeled cells. In order to quantify 4-Hydroxynonenal (4-HNE) levels in PVI free-floating sections were incubated in rabbit anti-parvalbumin (1:2000 working dilution; Swant Cat# PV 25, RRID:AB_10000344) and mouse anti-4HNE (1:1000 working dilution; Abcam Cat# ab48506, RRID:AB_867452) in PBS and 0.3% Triton X (Sigma-Aldrich) for 36 h at 4°C. Sections were washed three times for 10 min each in PBS before they were incubated in secondary 488 goat anti-rabbit (1:1000 working dilution; Jackson ImmunoResearch Labs Cat# 111-095-144, RRID:AB_2337978) and 647 goat anti-mouse (1:500 working dilution; Cell Signaling Technology Cat# 4410, RRID:AB_1904023) in PBS and 0.3% Triton X. To quantify 4HNE in PVI cells PVI immunofluorescence confocal images (20x magnification) from three sections of the prelimbic and infralimbic cortex were taken for each animal. ROIs were drawn around all PV+ cells in cellSens (Olympus cellSens Software, RRID:SCR_016238) and the mean gray intensity of the 4HNE signal for each cell was selected and averaged across each image and then across all three slices for every animal.

### GSH:GSSG Assay

In order to measure the ratio between reduced glutathione (GSH) and oxidized glutathione (GSSG), GSH, GSSG, and total glutathione were measured following the manufacturer instructions (glutathione detection kit, catalog #ADI-900-160, Enzo Life Sciences). In brief, animals were killed and the medial PFC containing the infralimbic and prelimbic cortex was dissected and homogenized in ice-cold 5% (w/v) meta-phosphoric acid (20 ml/g tissue), followed by centrifugation at 12,000 × g for 10 min at 4C. The resultant supernatant was collected for glutathione detection. For the measurement of GSSG and total glutathione, 2 M 4-vinylpyridine was added to the samples at a dilution of 1:50 (v/v). The samples were then incubated for 1 h at room temperature to derivatize reduced glutathione. Afterward, the samples were diluted in the reaction mix buffer. The reaction was observed by immediately and continuously recording changes at an optical density of 405 nm by using a microplate reader (Biotek) for a total of 15 min at 1 min intervals. The concentrations of total, oxidized, and reduced glutathione were normalized to the original wet weight of the tissue.

### Electrophysiology

Electrophysiological experiments used GFP+ hemizygous mice. Mice were anesthetized with urethane (3 g/kg body weight; Fisher Scientific) and transcardially perfused for one minute with gravity-fed ice-cold oxygenated (95% O2. 5% CO2) cutting ACSF, consisting of (in mM): 110 choline (Sigma-Aldrich), 25 NaHCO3 (Fisher Scientific), 1.25 NaH2PO4 (Fisher Scientific), 2.5 KCl (Sigma-Aldrich), 7 MgCl2 (Sigma-Aldrich), 0.5 CaCl2 (Sigma-Aldrich), 10 dextrose (Fisher Scientific), 1.3 L-ascorbic acid (Fisher Scientific), and 2.4 Na+-pyruvate (Sigma-Aldrich). Immediately after, brains were extracted and coronal sections (350 um) of the frontal cortex were cut on a vibratome (VT1000S, Leica) in cutting ACSF. Slices were transferred into a holding chamber containing warmed (35C) recording ACSF and cooled to room temperature over a one-hour period. The recording ACSF consisted of (in mM): 126 NaCl (Fisher Scientific), 25 NaHCO3, 1.25 NaH2PO4, 2.5 KCl, 2 MgCl2, 2 CaCl2, 10 dextrose, 2.4 Na+-pyruvate, and 1.3 L-ascorbic acid. For data collection, slices were transferred to a recording chamber affixed to an Olympus BX61WI microscope (Olympus) with continuous perfusion of oxygenated recording ACSF at room temperature. Whole-cell voltage-clamp recordings were obtained from pyramidal cells and PVIs in the prelimbic and infralimbic cortex using an Axon Multiclamp 700B amplifier (Molecular Devices). Data were acquired and analyzed using AxoGraph X (AxoGraph Scientific). Recording electrodes (WPI; 3–5 MΩ open tip resistance for pyramidal cells, 6–8 MΩ for interneurons) were filled with an internal solution consisting of (in mM): 130 CsCl (Sigma-Aldrich), 20 tetraethylammonium chloride (Sigma-Aldrich), 10 HEPES (Sigma-Aldrich), 2 MgCl2, 0.5 EGTA (Sigma-Aldrich), 4 Mg2+-ATP (Sigma-Aldrich), 0.3 Lithium-GTP (Sigma-Aldrich), 14 phosphocreatine (Sigma-Aldrich), and 2 QX-314 bromide (Tocris Bioscience). Theta-glass pipettes (Warner Instruments) connected to a stimulus isolator (WPI) were used for focal stimulation of synaptic potentials. Access resistance was monitored throughout the recording, and a <20% change was deemed acceptable. Spontaneous EPSCs were isolated by blocking chloride channels with the addition of picrotoxin (75uM; Sigma-Aldrich) into the recording ACSF. Spontaneous IPSCs were isolated by blocking AMPA receptor-mediated events with CNQX (6-cyano-7-nitroquinoxaline-2,3-dione; 20uM; Sigma-Aldrich). Miniature events were isolated by blocking sodium channels with the addition of tetrodotoxin (1uM; Alomone Labs). The frequency and amplitude of events were measured from 200 s of continuous recording using MiniAnalysis (Synaptosoft) with a threshold set at two times the RMS baseline noise. The ratio of currents through NMDA or AMPA receptors, respectively, was obtained by clamping cells at +40mv holding potential and applying local electrical stimulation. A compound evoked EPSC (eEPSC) was first recorded, then the AMPA component was isolated by washing CPP ((+/-)-3-(2-carboxypiperazin-4-yl)propyl-1-phosphonic acid; 10 uM; Sigma-Aldrich) into the bath. A minimum of 15 sweeps each were average for the compound and AMPA-only eEPSCs. The NMDA component was then obtained by digital subtraction of the AMPA component from the compound trace. The peak amplitude of the NMDA and AMPA traces were used to calculate the NMDAR/AMPAR ratio.

### Rule-Shifting

Procedures for our rule-shifting task followed those previously described (Phensy et al., 2017b). Mice were food restricted to 85% of their free-feeding weight over two weeks and handled for at least 5 minutes a day. Testing took place in white wooden plus maze (each arm is 10×34×15 cm, with a 10×10 cm center area) under low ambient illumination. The arms were labeled East, West, South, and North for reference. On days 4-6, the maze was converted into a T-maze by blocking off one of the arms with a divider, and additionally a visual cue (vertical black and white stripes on a 13×10 cm plastic sheet) was placed alternately near the entrance of one of the two choice arms in a pseudorandom manner (see below). During all days, reward pellets (Cheerio bits) were placed around the outside of the maze in order to prevent animals from using olfactory cues to infer the location of the reward. Mice were habituated to the maze over three days. On the first day of habituation, 4 reward pellets (1/8th Cheerios bits) were placed in each of the arms of the plus maze. Animals were placed into the center of the maze and were allowed to freely explore the maze for 15 minutes. If a mouse consumed all 16 pellets before the end of the habituation period, it was briefly placed in a holding cage while the maze was rebaited, and then the mouse was placed back into the maze until the end of the 15-minute period. On the second day of habituation, arms were baited with two pellets each, and on the third day of habituation only one food pellet was placed at the end of each arm. To reach habituation criterion, animals were required to consume all 4 food pellets at least 4 times within the 15-minute period. All animals in this study reached this criterion on the third habituation day. On the following day (Day 4), the plus maze was converted into a T-maze by blocking off one of the arms and the animals’ turn bias was determined. Therefore, mice were placed in the stem arm and allowed to turn left or right to obtain a food pellet. After the mouse consumed the reward, it was returned to the stem arm and allowed to make another choice. If the mouse chose the same arm as on the initial choice, it was returned to the stem arm until it chose the other arm and consumed the food pellet. Once both food pellets were consumed the maze was rebaited and the next trial began. The direction of the initial turn chosen four or more times over seven trials was considered the turn bias. On the next day (Day 5, Response Discrimination), mice were trained on an egocentric task which required them to always turn towards one side (left or right, chosen opposite to the direction of their turn bias) to obtain the food reward. The location of the stem arm was pseudorandomly rotated among 3 arms (East, West, and South) to discourage mice from using an allocentric spatial strategy. During all trials a visual cue was placed close to the entrance of one of the choice arms. Placement of this cue into the right or left arm varied pseudorandomly to balance the frequency of occurrences in each arm across blocks of 12 consecutive trials. Similarly, the order of the stem arms alternated pseudorandomly in a balanced fashion across blocks of 12 trials. Training continued until the mouse made 9 correct choices over 10 consecutive trials. When animals achieved this acquisition criterion, a probe trial was administered. In the probe trial the previously unused fourth arm (North) was used as a stem arm. If the mice performed the probe trial correctly, Response Discrimination training was completed. If an incorrect turn occurred, response training continued until the mouse made another five consecutive correct choices, and then another probe trial was administered. On the next day (Day 6, Shift-to-Visual-Cue Discrimination), mice were trained to shift their strategy to now select the choice arm with the visual cue in order to obtain food rewards. The location of the visual cue and the position of the start arm were again varied pseudorandomly so that their frequency was balanced across blocks of 12 consecutive trials. The training and response criteria for the Shift-to-Visual-Cue Discrimination were identical to those during Response Discrimination. Performance and Error Analysis: For each of the two test days we analyzed the total number of trials to criterion and the number of probe trials required to reach criterion. For the Shift-to-Visual-Cue Discrimination, errors were scored as entries into arms that did not contain the visual cue, and they were further broken down into three subcategories to determine whether the animals’ treatment altered the ability to either shift from the previously learned strategy (perseverative errors), or to maintain the new strategy after perseveration had ceased (regressive errors, or never-reinforced errors). In order to detect shifts in the strategies that animals used, trials were separated into consecutive blocks of four trials each. A perseverative error occurred when a mouse made the same egocentric response as required during the Response Discrimination, but which was opposite to the direction of the arm containing the visual cue. Six of every 12 consecutive trials required the mouse to respond in this manner. A perseverative error was scored when the mouse entered the incorrect arm on three or more trials per block of 4 trials. Once the mouse made less than three perseverative errors in a block, all subsequent errors of the same type were now scored as regressive errors (because at this point the mouse was following an alternative strategy at least half of the time). So-called never-reinforced errors were scored when a mouse entered the incorrect arm on trials where the visual cue was placed on the same side that the mouse had been trained to enter on the previous day.

### Novel Object Recognition

Testing was conducted in a white wooden open chamber (39 × 19 × 30.5 cm) and sessions were recorded from above by a web camera for later analysis. Wooden toys (approximately 3 × 5 cm) were used as stimulus objects and pseudorandomly selected as either the familiar or novel objects. In addition, in a different cohort of mice object preference was measured prior to experiments to ensure mice showed no inherent preference across the objects used. Mice were first habituated for 10 minutes on two consecutive days to the empty chamber. On the third day mice were again habituated for 10 minutes before the training and test trials begun. Therefore, mice were placed in their home cage while the chamber was cleaned and two objects were placed inside the chamber. Mice were then placed inside the chamber and allowed to investigate the two objects for 3 minutes before being placed back into the home cage for a 2-minute intertrial interval during which one of the two familiar objects was replaced with a novel object. After 2 minutes mice were placed back into the chamber and allowed to explore both the familiar and novel object for an additional 2 minutes. The objects were cleaned with 20% ethanol and the chamber was cleaned with 70% ethanol between animals. The amount of time the mice spent investigating the objects during both the training trial and the novel object trial were analyzed. In order to assess whether animals recognized the novel object as such we calculated a “recognition index”, which is the percentage of time spent investigating the novel object over the total investigation time for both objects.

### Social Interaction

Experimental mice and two size- and age-matched stimulus mice were housed individually for three days prior to the task. On the day of the test, the experimental mouse was placed into a new cage with 2.5 grams of their original bedding material to allow the animal to habituate for one hour. Stimulus mice were kept in a small custom cylindrical holding apparatus (height 20 cm, steel bars separated by 1 cm, acrylic base and lid), which could be placed inside the test cage. After one hour, the first stimulus mouse was placed in the holding apparatus and positioned into the cage with the test mouse for a trial interval of 1-minute while being recorded by an overhead camera. This was repeated for four trials with an intertrial interval of 10 minutes (Trials 1-4). On the fifth trial, a novel stimulus mouse was introduced into the cage to test for social recognition memory. All trials were recorded via an overhead camera and the interaction times (defined as sniffing and investigation of the stimulus mouse at close proximity) were analyzed for each trial.

### Statistical Analysis

Differences between groups were compared using one-way ANOVAs or two-way mixed ANOVAs as indicated. Post-hoc analyses using Tukey correction were used to determine specific group differences. All data is presented as mean ± standard error of the mean (SEM). An alpha level of p < 0.05 was considered significant.

## Results

### Perinatal KET-treatment reduces parvalbumin expression in adult mPFC in WT, but not in *Ppif*^*-/-*^ mice

A reduction in the number of parvalbumin-expressing interneurons (PVI) is a hallmark of schizophrenia (Lewis et al., 2005; Nakazawa et al., 2012) that is recapitulated by most animal models, including perinatal KET application (Jeevakumar et al., 2015; Phensy et al., 2017a). To determine if genetic deletion of Cyclophilin D can protect against KET-induced reductions in PVI, we performed immunohistochemistry and quantified the number of parvalbumin+ somata in the mPFC from both adult WT and *Ppif*^*-/-*^ mice which received either KET or saline during development (Fig. 1). A one-way ANOVA revealed a main effect of treatment on the number of PV+ cells over the number of DAPI+ cells (F_(3, 30)_ = 8.990, p < 0.001). Wildtype KET-treated mice showed a significant loss in PV expression. In contrast, *Ppif*^*-/-*^-KET mice were protected against KET-induced PVI loss and had similar numbers of PV+ cells than saline-control mice (Fig. 1B).

**Figure 1.**
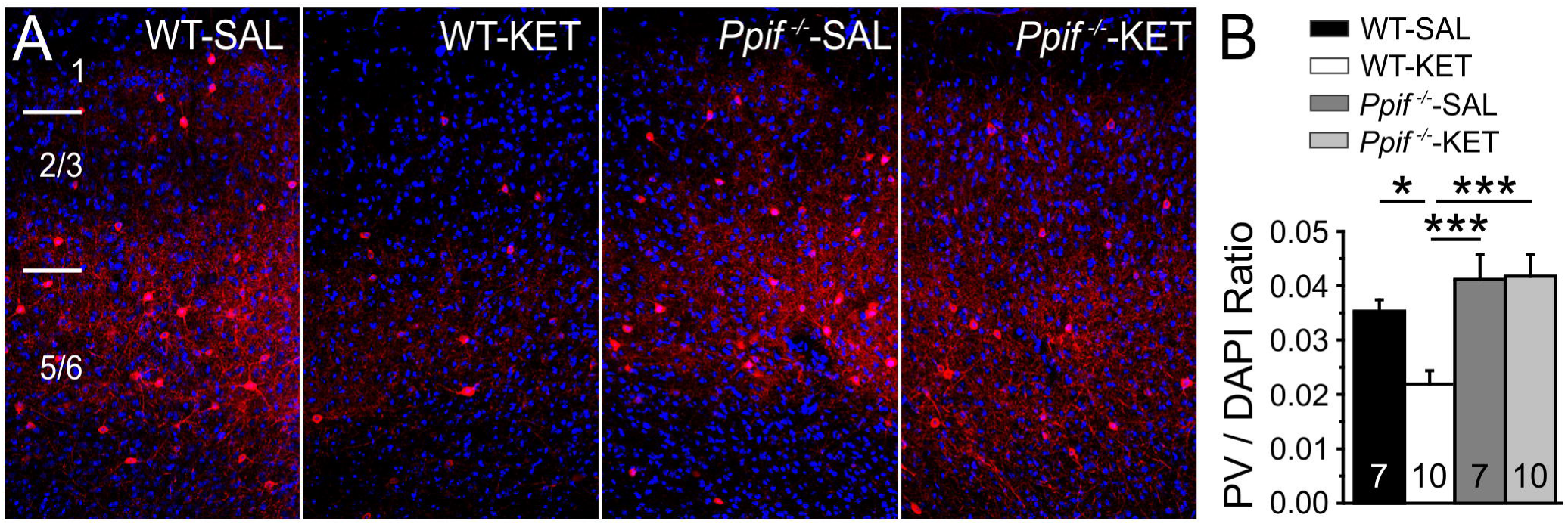
CypD-deletion (*Ppif*^-/-^) protects against the loss of parvalbumin (PV) expression that results from perinatal ketamine (KET) treatment. Wildtype (WT) or *Ppif*^*-/-*^ mice received either saline (SAL) or KET injections on postnatal days 7, 9, and 11. A) Representative confocal images of PV immunofluorescence (red) and DAPI (blue) from layers 1-6 of adult medial prefrontal cortex. B) Total PV-positive cells as a percentage of DAPI. Perinatal KET-treatment significantly reduced PV expression in WT but not in *Ppif*^-/-^ mice. Significance is indicated as *p ≤ 0.05, and ***p ≤ 0.001, following Tukey correction.

### Ketamine-induced oxidative stress is reduced in *Ppif*^*-/-*^ mice

Perinatal ketamine treatment leads to long-lasting increases in oxidative stress in adult animals, and PVI are particularly sensitive to redox dysregulation (Do et al., 2009; Phensy et al., 2017a). The ratio between the bioavailable reduced (GSH) and unavailable oxidized (GSSG) forms of the endogenous antioxidant glutathione provides a measure of redox balance in cells. A decrease in this ratio indicates a disruption in redox balance and subsequent oxidative stress. To determine if genetic deletion of cyclophilin D prevents KET-induced oxidative stress, we first measured levels of GSH and GSSG in mPFC tissue taken from adult mice (Fig. 2). A one-way ANOVA revealed that while there was no effect of treatment on total glutathione levels (F_(3, 14)_ = 1.172, p = 0.3556; Fig. 2A), there was a main effect of treatment on the ratio of GSH / GSSG (F_(3, 35)_ = 6.664, p = 0.001; Fig. 2B), with WT-KET mice having a significantly reduced GSH / GSSG ratio, indicating increased oxidative stress in these animals. A similar reduction was not observed in *Ppif*^*-/-*^-KET mice. Next, we measured 4-HNE levels in PVI of the mPFC (Fig. 3). 4-HNE levels increase during periods of oxidative stress due to lipid peroxidation. We colocalized immunofluorescence signals of 4-HNE and parvalbumin to measure changes in 4-HNE specifically in PVI. A one-way ANOVA revealed a main effect of treatment on the mean grey intensity of 4-HNE in PVI (F_(3, 27)_ = 5.084, p = 0.0064; Fig. 3B). This effect was due to a significant increase in 4-HNE signal in WT-KET mice, which was not present in *Ppif*^*-/-*^-KET or saline-control mice. Taken together, these results show that PVI in the mPFC of *Ppif*^*-/-*^ mice are protected from KET-induced oxidative stress.

**Figure 2.**
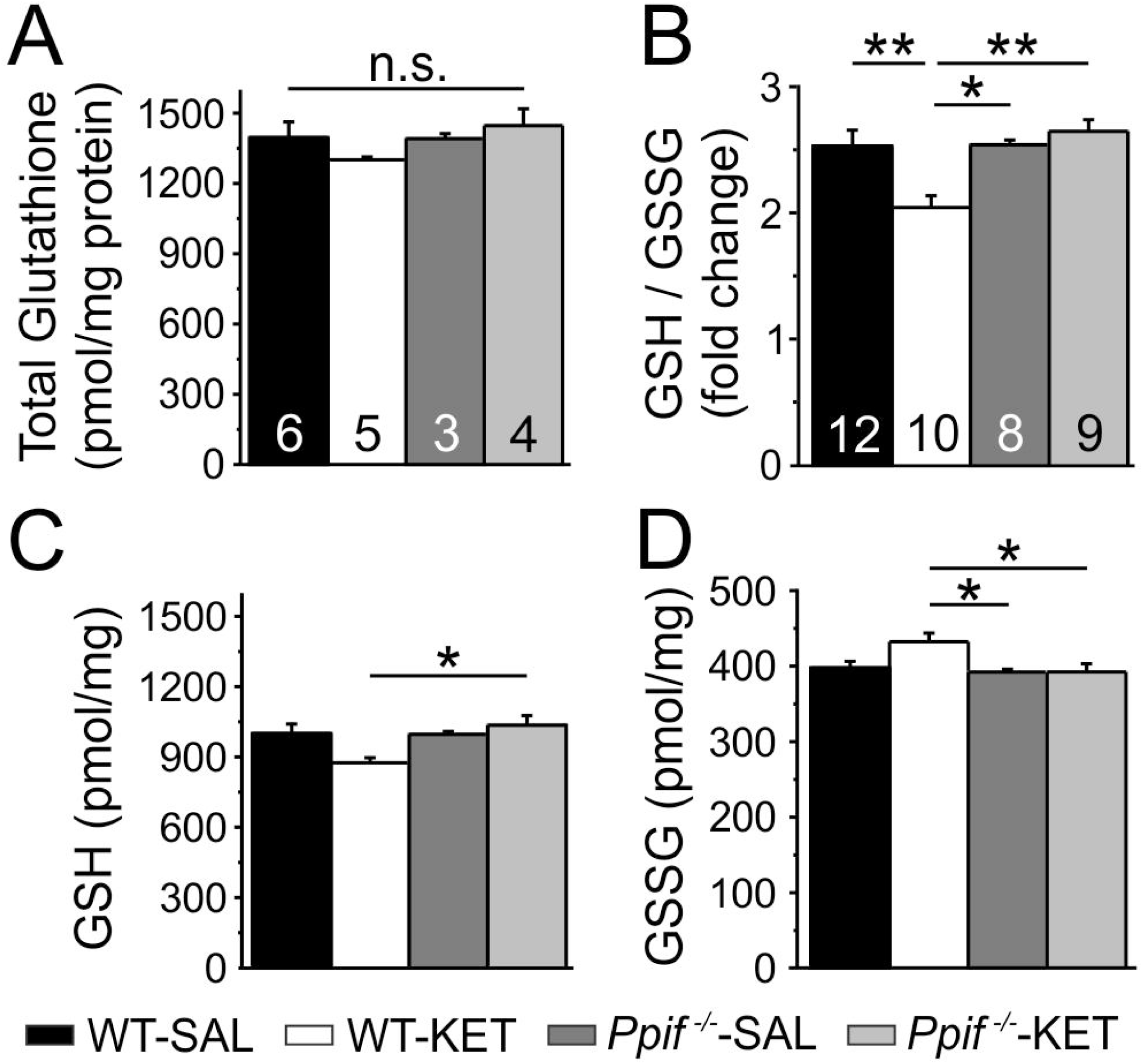
CypD-deletion (*Ppif*^-/-^) reduces perinatal ketamine (KET)-induced oxidative stress in adult medial prefrontal cortex (mPFC). A) KET-treatment does not affect total glutathione levels. B) KET-treatment significantly reduces the ratio of the reduced (GSH) over oxidized (GSSG) form of glutathione in wildtype (WT) mice, indicating increased oxidative stress. KET-treated *Ppif*^-/-^ mice are protected from this shift in the GSH / GSSG ratio. C, D) Total levels of GSH (C) and GSSG (D) in adult mPFC tissue. Significance is indicated as \**p* ≤ 0.05 and \*\**p* ≤ 0.01, following Tukey correction.

**Figure 3.**
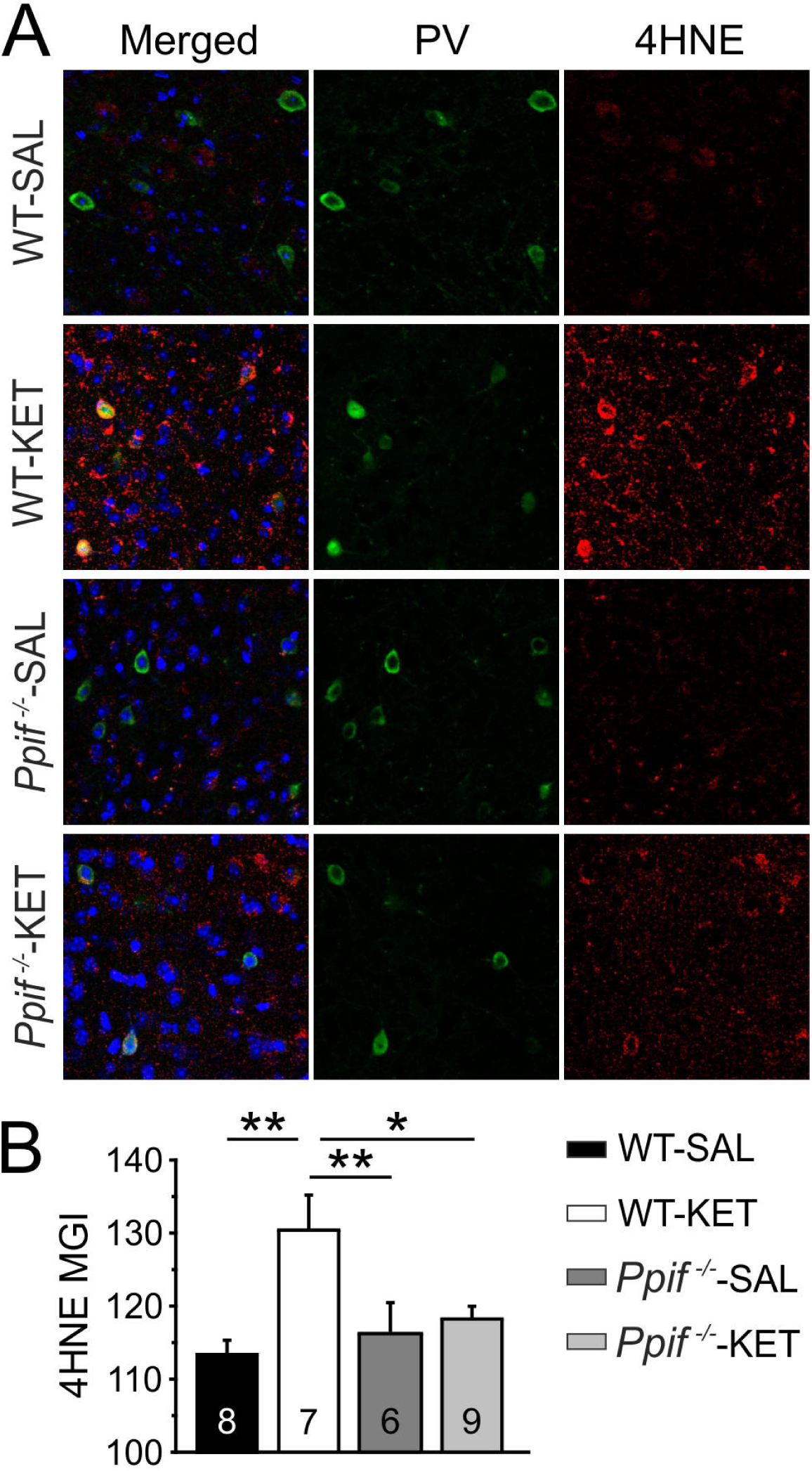
Perinatal ketamine (KET)-treatment increases oxidative stress in parvalbumin-positive interneurons (PVI) from wildtype (WT) but not from CypD knockout (*Ppif*^-/-^) mice. A) Representative confocal images of parvalbumin (PV, green) and 4-HNE (red) immunofluorescence in the medial prefrontal cortex. B) PV cells in tissue from WT-KET mice had significantly higher levels of 4HNE, a measure of lipid peroxidation and oxidative stress, than saline-treated WT mice (WT-SAL). KET-treated *Ppif*^-/-^ mice were protected from this increase. Significance is indicated as *p ≤ 0.05, and **p ≤ 0.01, following Tukey correction.

### Perinatal KET-treatment alters inhibitory synaptic transmission onto layer 2/3 mPFC pyramidal cells in WT, but not *Ppif*^*-/-*^ mice

GABAergic PVI inhibit nearby pyramidal neurons and regulate synchronized firing (Sohal and Rubenstein, 2019). Loss of PVI function leads to reduced GABAergic activity onto pyramidal neurons resulting in disinhibited circuits. In order to determine if CypD deletion prevents KET-induced disinhibition of pyramidal cells, we performed whole cell patch-clamp recordings in layer 2/3 pyramidal neurons of the mPFC and quantified the frequency and amplitude of both spontaneous (sIPSCs) and miniature inhibitory postsynaptic currents (mIPSCs) (Fig. 4). We found a significant main effect of treatment on the frequency of mIPSCs (F_(3, 28)_ = 6.547, p = 0.002; Fig. 4C), but no effect on amplitude (F_(3, 28)_ = 1.689, p = 0.192; Fig. 4C). Post-hoc analyses revealed that this was driven by a selective decrease in mIPSC frequency in WT-KET mice. Similar changes did not occur in *Ppif*^*-/-*^-KET or saline-control mice. KET-treatment did not alter the frequency (F_(3, 22)_ = 0.8743, p = 0.469; Fig. 4F) or the amplitude (F_(3, 22)_ = 1.289, p = 0.303; Fig. 4F) of sIPSCs. These data suggest that KET reduces GABA release and that this can be prevented by CypD deletion.

**Figure 4.**
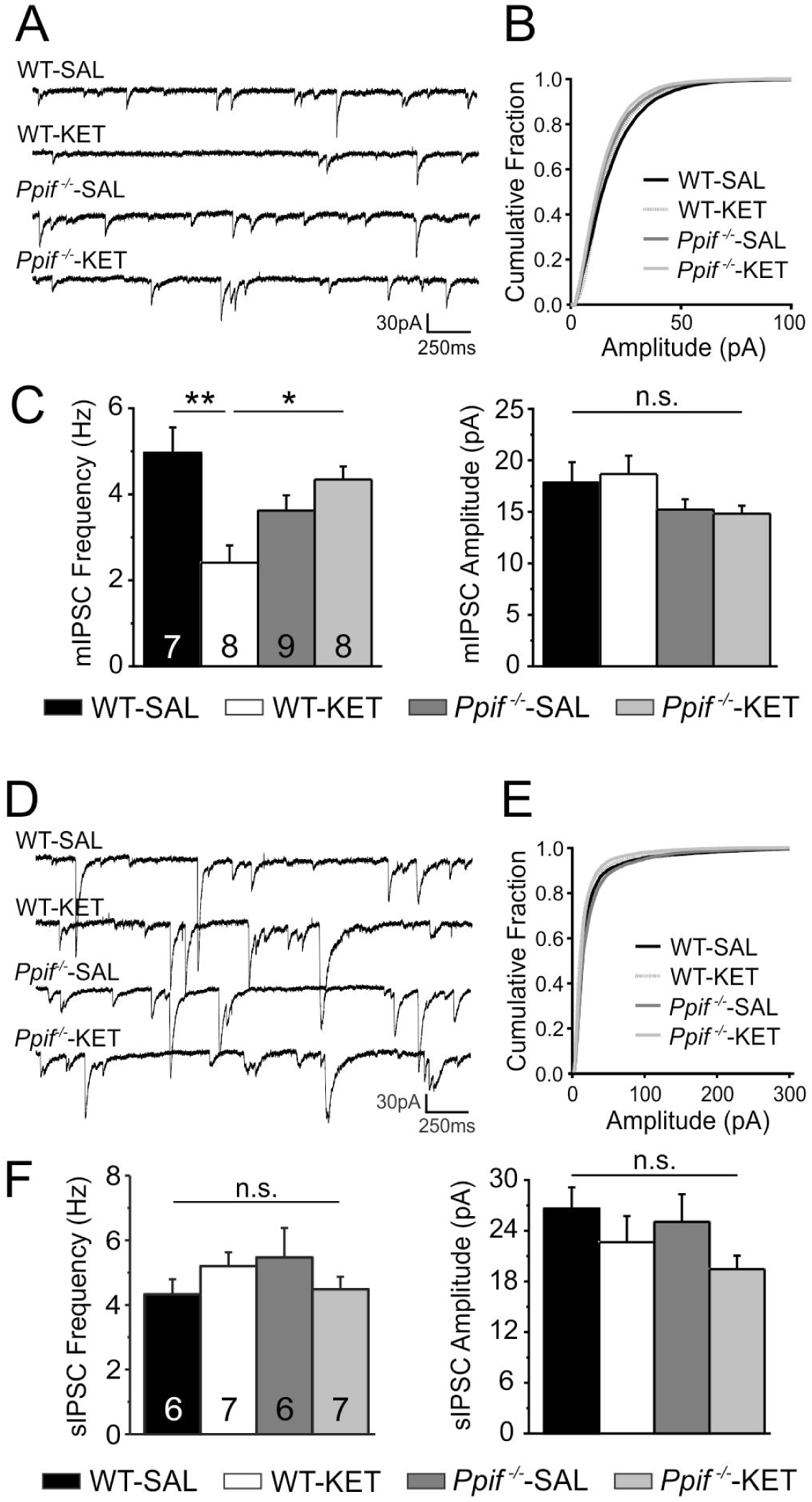
CypD deletion (*Ppif*^-/-^) protects against alterations in GABA release that develop in the medial prefrontal cortex following perinatal ketamine (KET)-treatment. A) Representative traces of miniature IPSCs (in tetrodotoxin) recorded in layer 2/3 pyramidal cells from in the 4 treatment groups. B) Amplitude distribution of all mIPSC events in the 4 treatment groups. C) KET-treatment significantly reduced the frequency, but not the amplitude of mIPSCs onto layer 2/3 pyramidal neurons from wildtype (WT) mice. Similar changes were not observed in *Ppif*^-/-^-KET mice. D) Representative traces of spontaneous IPSCs recorded in layer 2/3 pyramidal cells. E) Amplitude distribution of all sIPSC events in the 4 treatment groups. F) KET-treatment did not alter the frequency or amplitude of sIPSCs. Significance is indicated as *p ≤ 0.05, and **p ≤ 0.01, following Tukey correction.

### KET-treatment induces NMDAR hypofunction in layer 2/3 PVI from WT, but not *Ppif*^*-/-*^ mice

NMDAR hypofunction likely contributes to aberrant network activity in schizophrenia (Snyder and Gao, 2013). Perinatal KET treatment disrupts PVI development in the mPFC, causing NMDAR hypofunction in adult layer 2/3 PVI (Jeevakumar and Kroener, 2016; Phensy et al., 2017a). In order to test whether CypD deletion can prevent KET-induced changes in NMDAR-signaling we next measured NMDAR and AMPAR currents in GFP+ PVI (Fig. 5A-B). A one-way ANOVA (F_(3, 18)_ = 4.280, p = 0.019; Fig. 5B) revealed a main effect of treatment on the ratio of NMDAR:AMPAR currents at layer 2/3 PVI. Consistent with our previous reports (Jeevakumar and Kroener, 2016; Phensy et al., 2017a), WT-KET mice had significantly reduced NMDAR:AMPAR ratios. In contrast, *Ppif*^*-/-*^-KET mice showed current ratios comparable to saline-treated controls, suggesting that genetic deletion of CypD offers protection from KET-induced aberrant NMDAR signaling in layer 2/3 PVI.

**Figure 5.**
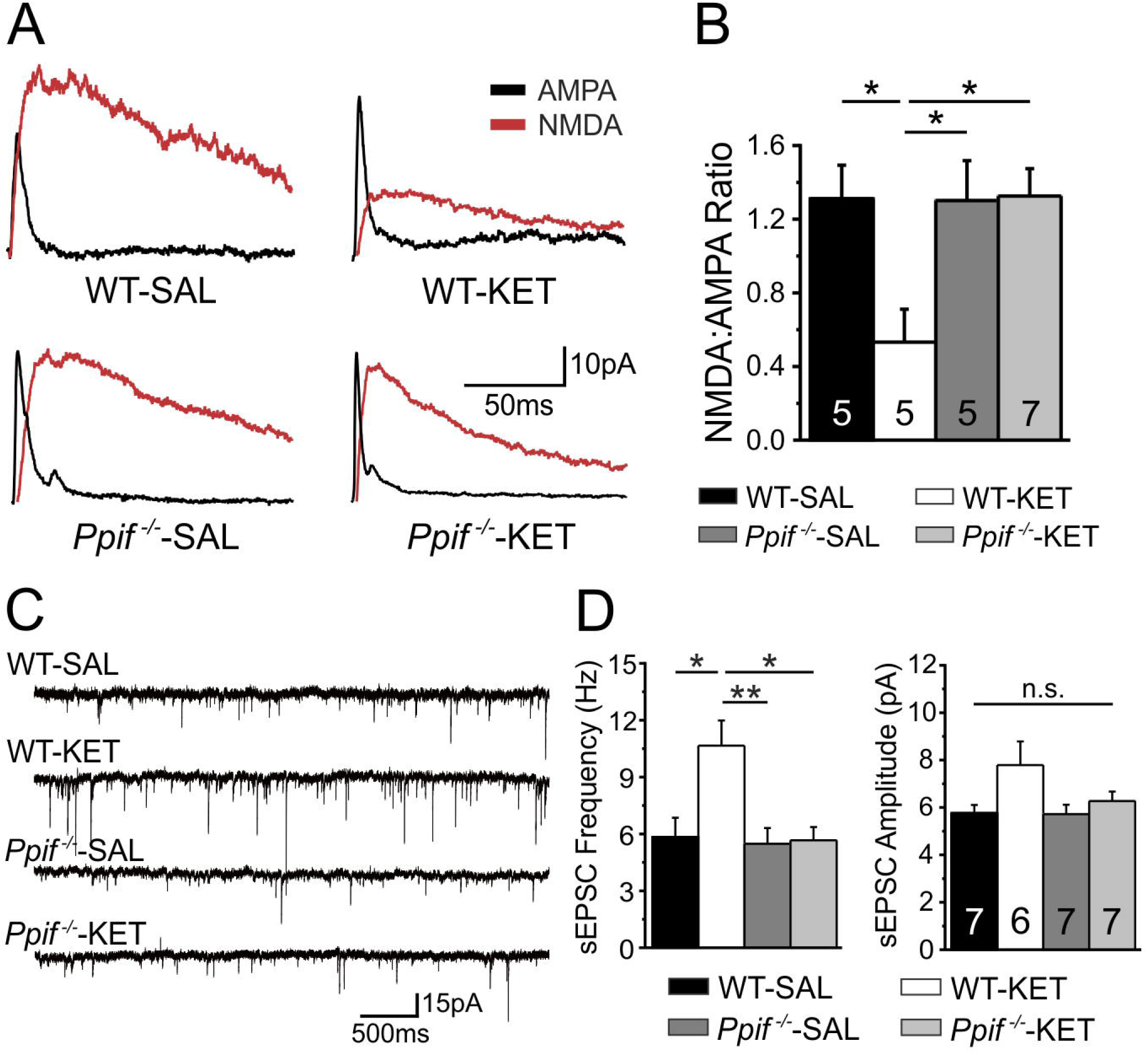
Perinatal ketamine (KET)-treatment alters glutamatergic transmission onto layer 2/3 parvalbumin-positive interneurons (PVI) in wildtype (WT), but not in CypD knockout (*Ppif*^-/-^) mice. A) Representative traces of NMDAR-currents (red) and AMPAR-(black) from GFP+ PVI neurons in layers 2/3 of the medial prefrontal cortex. B) NMDAR:AMPAR ratios in the 4 treatment groups. Ketamine-treated WT mice exhibit markedly decreased NMDAR:AMPAR ratios compared to saline (SAL)-treated mice, as well as KET-treated *Ppif*^-/-^ mice. C) Representative traces of spontaneous-EPSCs recorded at -70 mV. D) KET-treatment increased the frequency, but not the amplitude of sEPSCs in WT mice, but not in *Ppif*^-/-^ mice. Significance is indicated as *p ≤ 0.05, and **p ≤ 0.01, following Tukey correction.

### KET-treatment alters spontaneous glutamate release onto layer 2/3 PVI from WT, but not *Ppif*^*-/-*^ mice

The KET-induced NMDAR hypofunction in layer 2/3 PVI is accompanied by disinhibition of pyramidal cells (as seen in Fig. 4), which subsequently causes increased glutamate release back onto PVI (Jeevakumar and Kroener, 2016; Phensy et al., 2017a). The increased activation of postsynaptic glutamate receptors may lead to excessive calcium influx and contribute to persistent mitochondrial stress in PVI (Phensy et al., 2017a). To further test if CypD deletion prevents KET-induced alterations in glutamatergic signaling at PVI, we recorded spontaneous excitatory postsynaptic currents (sEPSCs) in GFP+ PVI (Fig. 5C-D). A one-way ANOVA revealed a main effect of treatment on sEPSC frequency (F_(3, 23)_ = 5.812, p = 0.004; Fig. 5D), without significant changes in sEPSC amplitude (F_(3, 20)_ = 2.828, p = 0.065; Fig. 5D). Consistent with the idea that mPFC pyramidal cells from KET-treated mice are disinhibited, post-hoc analyses showed a selective increase in sEPSC frequency in PVI from WT-KET mice; a change that was not seen in any of the other treatment groups.

### KET-treatment induces deficits in cognitive flexibility, novel object recognition, and social interactions in WT, but not in *Ppif*^*-/-*^ mice

In order to determine the functional impact of the physiological changes that result from KET-treatment and CypD deletion, we tested adult mice in a battery of behavioral tasks. These tasks included a rule-shifting task to measure cognitive flexibility, a novel object recognition task which measures (short-term) memory for objects, and a social interaction task which tests deficits in social interaction and novelty discrimination (Fig. 6) (Jeevakumar et al., 2015; Phensy et al., 2017b).

**Figure 6.**
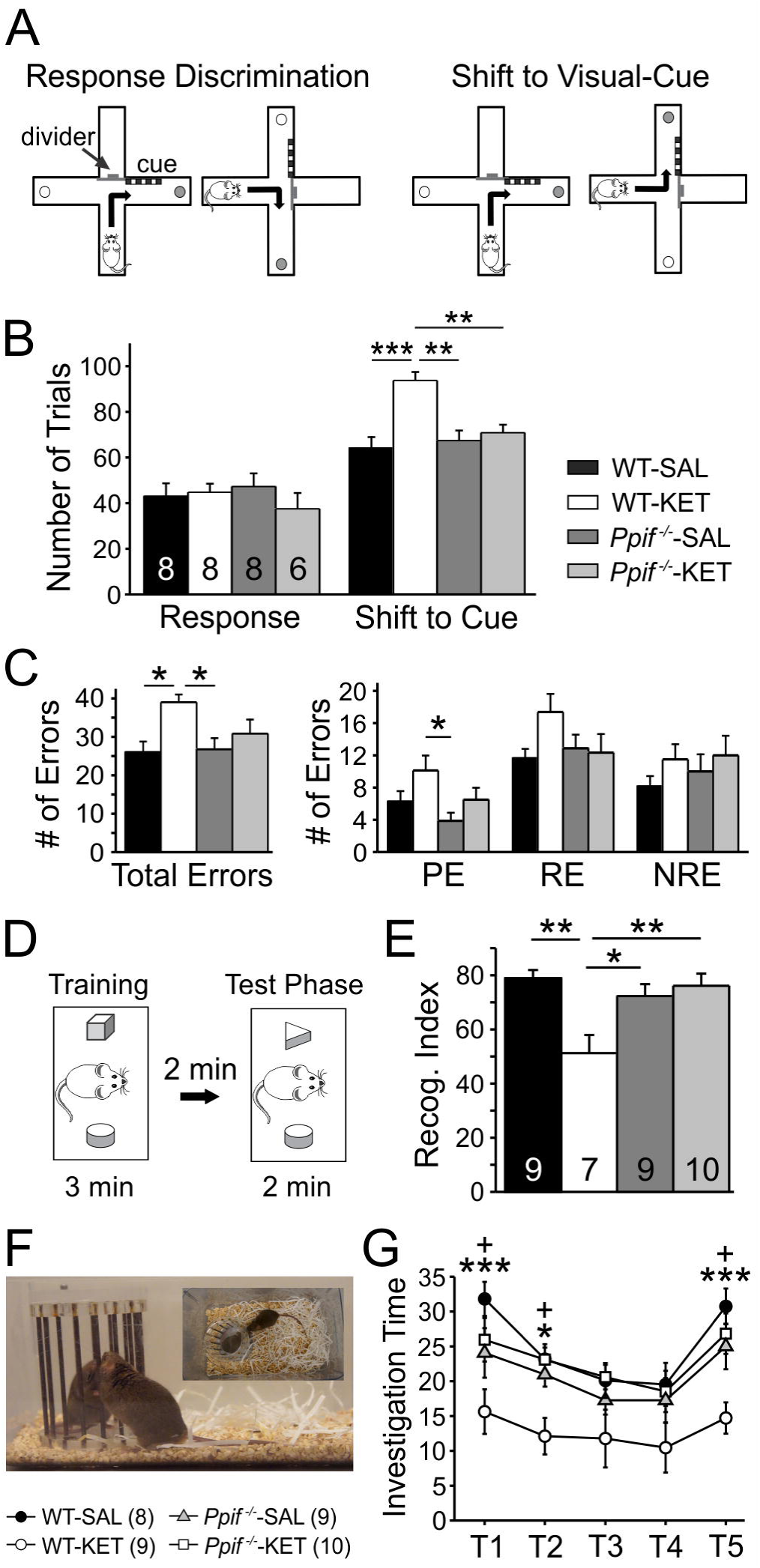
Perinatal ketamine (KET)-treatment induced behavioral deficits in wildtype (WT), but not in CypD knockout (*Ppif*^-/-^) mice. A) Schematic overview of the Cross-Maze Rule Shifting task used to assess cognitive flexibility. Mice learn an egocentric strategy (Response Discrimination) in order to obtain a food reward and the next day are required to shift to a visual cue strategy (Shift to Visual Cue Day). B) All animals learned the initial response strategy at the same rate; however, WT-KET required a significantly larger number of trials to shift between strategies compared to saline-treated controls and KET-treated *Ppif*^*-/-*^ mice. C) Error analysis based on error types committed during the Shift-to-Cue session. WT-KET animals show an increase in total errors and a significantly higher number of perseverative errors. D) Schematic overview of the setup used to test novel object recognition (NOR). E) KET-treatment reduced NOR in WT, but not in *Ppif*^*-/-*^ mice. F) Test of social interaction and recognition. A stimulus mouse is placed in the home-cage of the test mouse for four 1-minute sessions (10-minute inter-trial-intervals) and social interaction time is recorded. On a fifth trial, a novel stimulus mouse is introduced to test social recognition. G) KET-treatment results in reduced interaction times across all sessions. *Ppif*^*-/-*^-KET mice patterns of social interaction comparable to saline-treated controls. A-F: Significance is indicated as *p ≤ 0.05, **p ≤ 0.01, and ***p ≤ 0.001, following Tukey correction. G: Significance is indicated as*p ≤ 0.05 WT-SAL vs WT-KET, ***p ≤ 0.001 WT-SAL vs WT-KET, and +p ≤ 0.05 for WT-KET vs *Ppif*^*-/-*^-KET following Tukey correction.

### Rule Shifting Task

Cognitive flexibility is the ability to inhibit the use of a defunct strategy and enable the learning of a new functional strategy. The PFC is important for the ability to shift between strategies, and dysfunctions of the PFC lead to perseveration on inappropriate responses. To examine if CypD deletion protects against KET-induced deficits in cognitive flexibility, we tested WT and *Ppif*^*-/-*^ mice on a well-characterized rule-shifting task that is highly dependent on the mPFC (Birrell and Brown, 2000; Floresco and Magyar, 2006; Young et al., 2009; Hu et al., 2015; Jeevakumar and Kroener, 2016). Mice first learn an egocentric Response Discrimination strategy and then need to shift to a Visual-Cue Discrimination strategy. Mice in all treatment groups reached criterion for the Response Discrimination in the same number of trials (F_(3, 26)_ = 0.4760, p = 0.702; Fig. 6B). In contrast, a one-way ANOVA showed a main effect of treatment on the number of trials needed to reach criterion during the Shift-to-Visual Cue Discrimination phase of the task (F_(3, 26)_ = 9.576, p < 0.001; Fig. 6B). WT-KET mice took significantly more trials to reach criterion compared to mice in all other treatment groups. To further differentiate the effects of KET-treatment and CypD deletion on cognitive strategies, we analyzed the types of errors (perseverative, regressive, or never-reinforced) that mice committed. A one-way ANOVA (F_(3, 26)_ = 4.361, p = 0.013; Fig. 6C) revealed that KET-treatment caused significantly more overall errors in WT, but not *Ppif*^*-/-*^ or saline-control mice. Furthermore, there was a significant effect of treatment on perseverative errors (F_(3, 26_) = 3.152, p = 0.042; Fig. 6C), but not on regressive (F_(3,_ 26) = 1.867, p = 0.160; Fig. 6C), or never-reinforced errors (F_(3, 26)_ = 0.7296, p = 0.544; Fig. 6C). The significant effect on perseverative errors was due to a selective increase in the WT–KET mice which was not observed in KET-treated *Ppif*^*-/-*^ or saline-control mice.

### Novel Object Recognition

Mice, like humans, show preference for novel objects and spend more time investigating a novel object if they can correctly distinguish it from a previously encountered object. We next measured how KET treatment and CypD deletion affect novel object recognition (Fig. 6D-E). Preference for the novel object can be calculated as a recognition index, which is the time spent investigating the novel object as a percent of total time investigating both objects. A one-way ANOVA across treatment groups (F_(3, 31)_ = 6.375, p = 0.002; Fig. 6E) revealed a main effect of treatment on recognition index. WT-KET mice had reduced recognition indices; in contrast, *Ppif*^*-/-*^-KET exhibited normal novel object recognition and spent similar amounts of time as saline-controls investigating the novel object.

### Social Interaction

Deficits in social cognition and interaction greatly impact the quality of life of patients with schizophrenia. KET-treated mice show reduced social interactions (Jeevakumar et al., 2015; Phensy et al., 2017b). To test if genetic deletion of CypD can prevent this social deficit, we performed a social interaction and recognition task (Fig. 6F-G). Mice initially show great interest anytime a new mouse is introduced into their home cage but gradually reduce their investigation time with repeated exposures (trials 1-4). This can be used to investigate differences in baseline social interaction and recognition memory when a new stimulus mouse is introduced into the cage (trial 5). A two-way mixed ANOVA, revealed a significant main effect of treatment (F_(3, 32)_ = 7.954, p < 0.001; Fig. 6G) on social interaction. Consistent with our previous reports (Jeevakumar et al., 2015; Phensy et al., 2017b), WT-KET mice showed reduced investigation times across trials 1, 2, and 5. In contrast, *Ppif*^*-/-*^-KET mice demonstrated investigation times similar to saline-treated controls across all five exposures, suggesting normal social interaction and recognition memory.

## Discussion

NMDAR dysfunction disrupts normal development of GABAergic and glutamatergic networks and this may contribute to schizophrenia pathology (Krystal et al., 2002). Parvalbumin-expressing interneurons appear to be particularly susceptible to NMDAR dysfunction (Cohen et al., 2015), and changes in PVI and their synapses are well-documented in schizophrenia (Lewis et al., 2005; Nakazawa et al., 2012; Gonzalez-Burgos et al., 2015). PVI are highly sensitive to oxidative stress (Do et al., 2009), which can disrupt neuronal function and decrease NMDAR activity (Choi and Lipton, 2000). Mitochondria are the primary mediators of redox state (Rego and Oliveira, 2003; Bhatti et al., 2017) and they are abundant in PVI (Gulyas et al., 2006); however, their role in PVI dysfunction has received relatively little attention. Oxidative and other cellular stresses trigger translocation of CypD to the inner mitochondrial membrane, initiating formation of the mPTP (Baines et al., 2005). Prolonged mPTP formation leads to excessive levels of intracellular superoxide and pathological mitochondrial activity (Crompton, 2004; Lemasters et al., 2009). Glutamate excitotoxicity that results from overactivation of NMDA receptors and excessive calcium entry is a well-established initiator of chronic mPTP formation (Schinder et al., 1996).

NMDAR blockade disrupts development of PVI, alters E/I balance, and impairs cognitive performance (Wang et al., 2008; Jeevakumar and Kroener, 2016). There is strong evidence that these changes are mediated by oxidative stress (Radonjic et al., 2010; Powell et al., 2012), and we previously demonstrated that boosting antioxidant defense systems with N-acetyl cysteine can counter the physiological and behavioral deficits induced by perinatal KET-treatment (Phensy et al., 2017a). We also found that perinatal KET-treatment significantly increased levels of mitochondrial-derived ROS and reduced mitochondrial membrane potentials in PVI. These changes are signs of prolonged mPTP activation, suggesting mitochondria as important nodes in KET-induced PVI dysfunction. CypD is a necessary component of the mPTP, and reducing CypD translocation protects mitochondrial function (Du and Yan, 2010). Thus, we hypothesized that *Ppif*^*-/-*^ mice would be protected from KET-induced increases in oxidative stress and PVI dysfunction.

We measured glutathione levels to determine the redox state of PFC tissue from adult KET- and SAL-treated mice. The ratio of GSH to GSSG indicates cell redox status, with healthy cells having a large GSH/GSSG ratio, that drops when they get exposed to oxidative stress (Pizzorno, 2014). Wildtype KET-treated mice showed a significant decrease in the GSH/GSSG ratio in PFC. This is in line with previous findings by us (Phensy et al., 2017a) and others (Powell et al., 2012) which have shown that NMDAR blockade during development drives oxidative stress in the frontal cortex. We also found increased levels of 4-HNE, a byproduct of lipid peroxidation, in prefrontal PVI of WT-KET mice. Lipid peroxidation occurs when free radicals damage lipids and it is an indicator of the damage that results from oxidative stress. *Ppif*^*-/-*^-KET mice demonstrated robust protection against KET-induced reductions in the GSH/GSSG ratio and the increase of 4-HNE levels in PVI (Figs 2,3). Because *Ppif*^*-/-*^-KET mice also showed no significant loss of PV immunofluorescence in the PFC these results support the idea that PVI dysfunction results from mitochondrial oxidative stress. Previous reports have shown that PVI dysfunction following perinatal NMDAR blockade requires activation of NADPH-oxidase 2 (NOX2) (Behrens et al., 2007; Sorce et al., 2010). Interestingly, there is evidence for significant crosstalk between mitochondria and NOX2, which can reciprocally drive ROS production (Dikalov, 2011; Daiber et al., 2017). Thus, our data support these studies and suggest a complementary mechanism to NOX2-mediated PVI deficits.

NMDAR hypofunction in PVI is believed to contribute to aberrant synaptic activity in schizophrenia (Homayoun and Moghaddam, 2007). NMDARs can be directly inhibited by oxidizing agents via interaction on a redox-sensitive site on the receptor (Choi and Lipton, 2000). Developmental NMDAR blockade with ketamine leads to both long-lasting increases in oxidative stress and NMDAR-hypofunction in layer 2/3 prefrontal PVI (Phensy et al., 2017a). Because *Ppif*^*-/-*^-KET mice exhibited reduced signs of oxidative stress, we investigated if this was accompanied by normal NMDAR function in layer 2/3 PVI. Consistent with our previous findings (Jeevakumar and Kroener, 2016; Phensy et al., 2017a), PVI in layers 2/3 from KET-treated WT mice showed reduced NMDAR currents. In contrast, *Ppif*^*-/-*^-KET mice exhibited normal NMDAR:AMPAR current ratios (Figure 4). Because KET-treatment did not affect amplitudes of AMPA-mediated sEPSCs, this effect of CypD-deletion most likely represents a selective protection of NMDAR function. Reduced NMDAR activity has significant implications for PVI function. Gating of NMDARs causes influx of calcium which can help in persistent neuronal firing (Myme et al., 2003), an important feature of PVI physiology. Blocking NMDAR on PVI has been shown to impair the generation of gamma oscillations (Jadi et al., 2016), which are crucial to cognitive function (Fries, 2009; Sohal et al., 2009). Perturbations in gamma oscillations are believed to result from reduced PVI activity and a shift in the excitation/inhibition (E/I) balance (Gonzalez-Burgos et al., 2015; Sohal and Rubenstein, 2019). Consistent with the idea of a shift in the E/I balance, KET-treatment in WT mice lead to a long-lasting reduction in GABAergic inhibition at pyramidal neurons and increased glutamate release back onto layer 2/3 PVI. In contrast, in addition to preserved NMDAR function in PVI, *Ppif*^*-/-*^ mice exhibited normal sEPSCs and mIPSCs in PVI and pyramidal cells, respectively, suggesting that normal E/I balance in the mPFC network was maintained.

Patients with schizophrenia suffer from a number of PFC-dependent cognitive deficits including disruptions in working memory, social cognition, attention, and cognitive flexibility (Braff et al., 1991; Gold et al., 1997; Nuechterlein et al., 2004). Evidence from clinical and preclinical models suggests that NMDAR hypofunction contributes to these deficits (Coyle, 2012; Cohen et al., 2015): NMDAR blockade can reduce cognitive abilities in healthy patients and exacerbate deficits in schizophrenia patients (Lahti et al., 1995; Malhotra et al., 1997; Krystal et al., 2002), and it impairs cognitive flexibility, episodic memory, and social interactions in rodents (Stefani and Moghaddam, 2005; Powell et al., 2012; Jeevakumar et al., 2015). In order to determine if CypD-deletion also protects against KET-induced cognitive deficits we examined the performance of *Ppif*^*-/-*^-KET mice in a variety of tasks (Figure 6). In rodents, cognitive flexibility is most often assessed via attentional set-shifting tasks (Young et al., 2012). Here, we measured cognitive flexibility through a rule-shifting task which requires only a simple shift from an egocentric response strategy to a visual cue-based strategy (Stefani and Moghaddam, 2005; Floresco et al., 2006). Consistent with previous findings (Stefani and Moghaddam, 2005; Broberg et al., 2008; Jeevakumar et al., 2015) we found that WT-KET mice required more trials to shift their strategies and committed a larger number of perseverative errors (Figure 6A-C). Perseverative errors suggest an inability to abandon a defunct strategy, a deficit that is frequently observed in patients with schizophrenia (Abbruzzese et al., 1996) or lesions of the PFC (Barcelo and Knight, 2002). Patients with schizophrenia also suffer from deficits in episodic memory (Ragland et al., 2009). In rodents, episodic memory can be assessed via the novel object recognition task. Performance on the task relies heavily on interactions between PFC and hippocampal circuits (Korotkova et al., 2010), which are disrupted by blockade (Jadi et al., 2016) or ablation (Korotkova et al., 2010) of NMDARs. Both acute (Rajagopal et al., 2014) and developmental (Jeevakumar et al., 2015; Phensy et al., 2017b) KET-treatment results in reduced novel object recognition. Finally, we examined changes in social interaction in KET-treated WT and *Ppif*^*-/-*^ animals. Reduced social interactions and isolation are negative symptoms associated with schizophrenia (Millan et al., 2014; Green et al., 2015). Consistent with what we (Phensy et al., 2017b) and others (Powell et al., 2012) have previously shown, developmental NMDAR blockade in WT mice reduced social interaction times across all presentations of the stimulus mice. Importantly, *Ppif*^*-/-*^ mice showed robust protection against all KET-induced behavioral deficits. These findings strongly suggest that transient NMDAR blockade affects cortical networks and behavior via processes that depend on proper mitochondrial function, and that modulation of the mPTP via genetic deletion of CypD can prevent these effects. These findings are in line with a number of other studies in which genetic deletion or pharmacological inhibition of CypD has been shown to offer protection against cognitive dysfunction in other preclinical disease models (Du et al., 2008; Yan et al., 2016; Nusrat et al., 2018). One previous study reported higher indices of anxiety and a reduced tendency to explore in *Ppif*^*-/-*^ mice (Luvisetto et al., 2008); however, we did not find evidence for reduced exploration during NOR or the cross-maze rule-shifting task, nor did we observe any other unspecific phenotypical changes in *Ppif*^*-/-*^ mice.

Taken together, our results underscore the impact of mitochondria on cortical networks and cognition. Mitochondria not only play essential roles in cell function, but their bioenergetics are crucial for proper neuronal development (Cobley, 2018), and even acute dysfunction impairs learning and memory (Mancini and Horvath, 2017). Here we illustrate how CypD activity can drive mitochondrial dysfunction in PVI and show that CypD may be a potential therapeutic target in protecting cognitive function in schizophrenia.

